# Repellent compound with larger protective zone than DEET identified through activity-screening of *Ir40a* neurons, does not require *Or* function

**DOI:** 10.1101/017145

**Authors:** Tom Guda, Pinky Kain, Kavita Sharma, Christine Krause Pham, Anandasankar Ray

**Affiliations:** Department of Entomology, Center for Disease Vector Research, 3401 Watkins Drive, Riverside, CA.; Interdepartmental Neuroscience Program, University of California Riverside, 3401 Watkins Drive, Riverside, CA.

**Keywords:** Insect repellents, DEET, *Ir40a*, calcium imaging, mosquito, *Orco*

## Abstract

The widely used insect repellent DEET has a limited spatial zone of protection, requiring it to be applied over all exposed areas of skin. Identification of insect DEET-sensing neurons expressing a highly conserved *Ionotropic receptor*, *Ir40a*, provides an opportunity to identify new structural classes of volatile agonists as potential spatial repellents. By imaging the activity of the *Ir40a+* neurons in *D. melanogaster* expressing the calcium sensitive GCaMP3 protein, we identify a strong agonist, 4-methylpiperidine, with a much higher vapor pressure than DEET. Behavioral testing reveals that 4-methylpiperidine repels *Aedes aegypti*, which is consistent with our model that *Ir40a* marks a conserved innate aversive pathway. Using a spatial repellency assay we demonstrate that 4-methylpiperidine applied to one part of the hand repels mosquitoes on another part effectively, whereas DEET cannot do so. Using *orco* mutant *A. aegypti* we demonstrate that avoidance to 4- methypiperidine is not dependent on *Or* family function. Additional testing of *orco* mutant mosquitoes demonstrates that they are also effectively repelled by DEET, without coming in contact with it, in heat attraction assays. Together, these results support our initial observations that the conserved *Ir* pathway plays a key role in olfactory repellency and can be used to identify new classes of repellents.

## Introduction

Mosquitoes and other blood-feeding insects transmit deadly diseases such as malaria, dengue, lymphatic filariasis, West Nile fever, Yellow fever, sleeping sickness and Leishmaniasis to hundreds of millions of people, causing severe suffering and more than a million deaths each year. Current methods such as insecticide treated bed nets and indoor residual sprays provide the primary line of protection against malaria transmission. However, the use of volatile attractants and/or repellents may provide additional lines of defense and diminish concerns about effects on human health and emergence of resistance in mosquitoes, which are associated with heavy insecticide use.

*N,N*-Diethyl-m-toluamide (DEET) is one of the few compounds known to elicit aversion across most species of insects and arthropods and has remained the most popular insect repellent for over 60 years. However it plays a very limited role in the global context due to its relative cost, and the inconvenience of requiring continuous application to exposed skin at high concentrations. DEET has been shown to weakly inhibit human acetylcholinesterase (Corbel, Stankiewicz et al. 2009; Swale, Sun et al. 2014), and also block mammalian Na+ and K+ ion channels, which may be contributing factors in numbness that occurs if DEET is applied to the lip (Swale, Sun et al. 2014). Several instances of increased resistance to DEET have also been reported in flies (Reeder, Ganz et al. 2001) as well as mosquitoes (Klun, Strickman et al. 2004; Stanczyk, Brookfield et al. 2010). Furthermore, DEET is a strong plasticizer capable of dissolving plastics, synthetic fabrics, and painted and varnished surfaces (Krajick 2006), precluding its use in bed nets and in many urban settings.

Recent studies have put forth competing models regarding the mechanisms of DEET action (Ditzen, Pellegrino et al. 2008; Syed and Leal 2008; Xia, Wang et al. 2008; Liu, Pitts et al. 2010; Stanczyk, Brookfield et al. 2010; Pellegrino, Steinbach et al. 2011; Bohbot and Dickens 2012; Xu, Choo et al. 2014). Our own efforts to elucidate mechanisms of DEET repellency have identified *Ir40a* as a DEET-detecting receptor expressed in neurons innervating the sacculus of the *Drosophila melanogaster* olfactory system (Kain, Boyle et al. 2013). Identification of these DEET-responsive neurons in antennae of the genetically tractable *Drosophila melanogaster* provides an excellent assay system to identify new classes of broad-spectrum insect repellents.

We recently developed a cheminformatics descriptor-based prediction algorithm to screen >400,000 compounds (including 3000 natural compounds) on the basis of structural similarity to DEET and related repellents, and identified numerous candidate repellents. Approximately 80% of the predicted repellents activates *Ir40a+* neurons and were strong repellents in insects (Boyle, McInally et al. 2013; Kain, Boyle et al. 2013; Afify, Horlacher et al. 2014). Many of these chemicals do not dissolve plastic, are affordable, have pleasant odors, are of natural origin and have safer oral/dermal toxicity values as compared to DEET. Such efforts to identify new repellents have traditionally involved testing of structural analogs of DEET, but they do not allow for identification of structurally diverse chemical classes of repellents.

## Results

### Discovery of a structurally distinct activator of the Ir40a+ neuron using imaging

We wanted to test whether the newly described DEET-detecting *Ir40a+* neurons in *Drosophila* can help identify new chemical classes of ligands as repellents. Using the *GAL4/UAS* system, we expressed the calcium indicator GCaMP3 in *Ir40a*+ neurons and screened compounds for their ability to activate these neurons using calcium imaging. This approach identified 4-methylpiperidine as an agonist of Ir40a neurons. More rigorous testing revealed that a 2-sec stimulus delivered from a filter paper spotted with 20% 4-methylpiperidine activated *Ir40a* neurons in a sustained manner for more than 3 minutes, while the water solvent alone did not (Fig 1a,b). The prolonged temporal kinetics of the response was similar to what was previously observed for DEET and other ligands (Kain, Boyle et al. 2013), and is consistent with the notion that molecules entering the sack-like sacculus would circulate inside it longer due to restricted air movement, unlike on the antennal surface. The level of activation measured as %ΔF/F for 4-methylpiperidine was >700%, with none observed for solvent controls (water and paraffin oil) or other odors at same concentration (Fig 1c). The activation level of this agonist was dramatically greater than previously observed for DEET and related compounds (∼60%-300%) (Kain, Boyle et al. 2013). Comparison of the chemical structure of 4-methylpiperidine show very little structural overlap with DEET and four other computationally identified agonists of *Ir40a* neuron that are repellents (Kain, Boyle et al. 2013). The vapor pressure of this smaller agonist is substantially higher than that of previously identified agonists of the *Ir40a* neuron, which could perhaps contribute to its stronger activation (Fig 1d).

**Figure 1.**
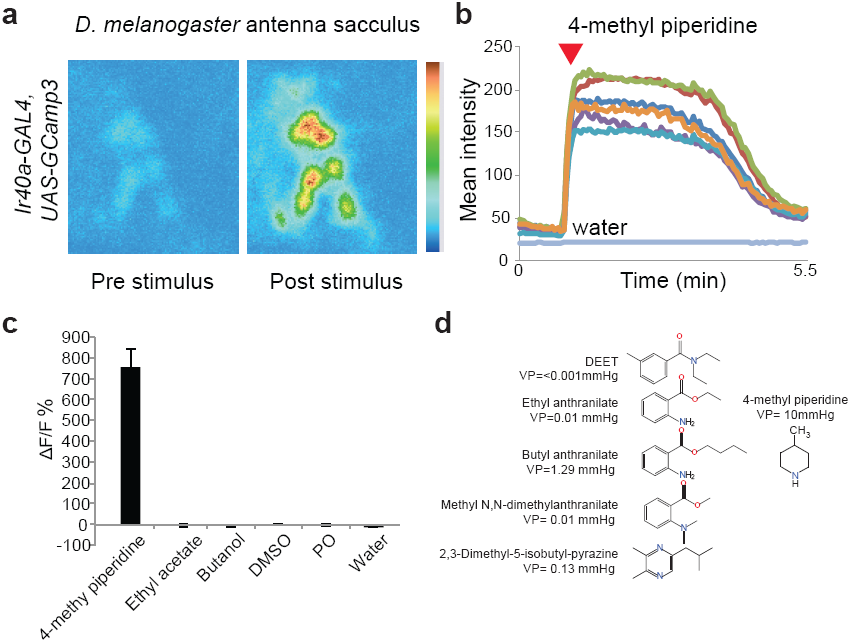
Identification of a strong *Ir40a+* neuron agonist with structure unrelated to DEET and increased volatility. Odor-evoked calcium responses from the sacculus neurons of individual antenna of *Ir40a- Gal4/UAS-GCaMP3* flies. **a,** Representative fluorescence image of the antennal sacculus neurons expressing GCaMP3 with color-coded heat map representing changes in fluorescence intensity before and after a 2-sec puff of 4-methylpiperidine (4MPD) at 10%. **b**, Graph depicting changes in mean fluorescent intensity for 6 different cells (lines in different color) in the antenna in response to a ∼2-sec puff of 4MPD (red arrowhead) or the water solvent. **c,** Mean percentage change in fluorescent intensity ratio after a 2-sec odor stimulus as indicated N=9-18, error bars =s.e.m. Representative structures of DEET and four other previously identified agonists of the neuron alongside 4MPD with vapour pressure values indicated in parenthesis.

### Ir40a-neuron agonist repels flies and mosquitoes

We have previously shown that agonists of the *Ir40a-*neuron were all repellents when tested in behavioral assays in flies and in mosquitoes (Kain, Boyle et al. 2013). However, these chemicals were all identified by structural similarity to DEET and did not rule out the small possibility that the repellent effect of the four new compounds may be conveyed by another olfactory detection pathway, such as a specific *Or* family receptor. To test whether a *Ir40a* neuron agonist, which is structurally distinct from DEET, can also be repellent, we tested *D. melanogaster* in a T-maze behavior assay. We found that DEET is a weak repellent in this assay, as has been reported earlier, however the stronger agonist, 4-methylpiperidine was more than twice as aversive (Fig 2a). The finding that a structurally distinct agonist of the *Ir40a* neuron also imparted aversive valence supports our model *Ir40a* neurons mark a dedicated avoidance pathway.

**Figure 2.**
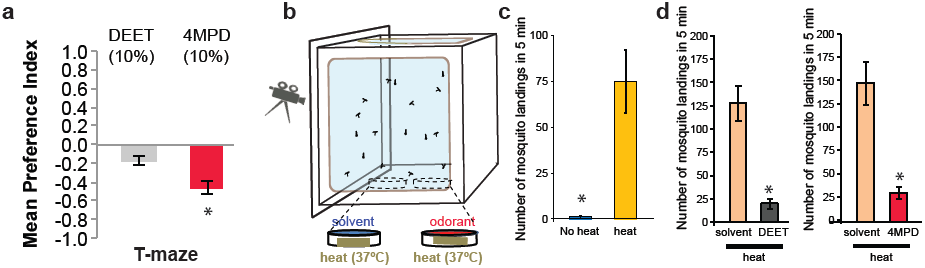
Ir40a+neuron agonist 4-methylpiperidine repels *Aedes aegypti* mosquitoes from an attractive heat source. **a,** Mean Preference index for indicated odorants in a T-maze with *Drosophila melanogaster,* N=10 trails each chemical, ∼40 flies/trial. Error bars= s.e.m. **b,** Schematic of the 2-choice heat attraction assay where two Petri dishes each with a ∼37°C heat source and a net-covering which is treated with test chemicals, is placed inside a cage containing 20 female *A. aegypti*. **c,** Total Mean number of mosquito landings counted in a 5 min period inside the cage, over the area of the two Petridishes. N=13 trials, 20 mosquitoes/trial. Error bars= s.e.m. **d,** Mean number of mosquito landings recorded in the area over each Petridish in pairs of 2-choice treatments. N=8 trials, 20 mosquitoes/trial. Error bars=s.e.m. ., *=p<0.005.

The *Ir40a* neuron in *D. melanogaster* co-expresses two additional members of the *Ionotropic receptor* family, *Ir93a* and *Ir25a*, all of which are well conserved in mosquitoes and other insects. The subunit composition of the receptor that responds to 4-methylpiperidine, or for that matter DEET, is not known yet, nor is their exact expression patterns in the mosquito olfactory organs. However, the high degree of amino acid conservation for all 3 Irs suggests that mosquitoes may also detect repellent ligands of the *Ir40a* neuron identified in *Drosophila*. In order to test behavioral conservation we tested 4-methylpiperidine for aversion in female *Aedes aegypti* mosquitoes. Female *A. aegypti* are attracted to warmth from the human body, which can be recapitulated in the laboratory using heat sources placed in the cages (Fig 2b,c). When DEET was added to a net placed above one of the two heat sources in a 2-choice paradigm, mosquitoes strongly avoided the heat source with the treated net (Fig 2c,d). The results of this assay indicate that mosquitoes detect DEET, in the absence of other odorant cues, and its presence causes avoidance of an otherwise attractive heat source. 4-methylpiperidine tested in this assay showed a strong repellent effect as well (Fig 2d). The finding that a structurally distinct agonist of the *Ir40a-*neuron evokes similar aversive behavior suggests that this is an olfactory avoidance pathway that is conserved.

### Spatial repellency imparted by volatile 4-methylpiperidine

One of the drawbacks of DEET is its limited spatial range of repellent activity, which necessitates its application across every square inch of exposed skin to protect against mosquito bites. Since 4- methylpiperidine is more volatile than DEET, we reasoned that its repellent effect might extend over a larger spatial zone. We performed experiments to test this possibility. We used a modified arm-in-cage experiment to test mosquitoes, in which we applied DEET or 4-methylpiperidine to filter paper surrounding the net-covered window that gave access to an area above the skin of a human hand inserted in a glove. When DEET (3%) was applied, mosquitoes were not repelled from the net-covered window, demonstrating that the effectiveness of DEET does not extend to the 1 cm border (Fig 3a,b). By contrast, application of 4-methypiperidine (3%) to the filter paper was extremely effective in repelling mosquitoes from the net window (Fig 3a,c). Although 4-methylpiperidine is not safe for human skin application, these proof-of-principle experiments indicate that identification of new classes of *Ir40a-*neuron activators can lead to development of novel repellents with substantially improved spatial protection properties.

**Figure 3.**
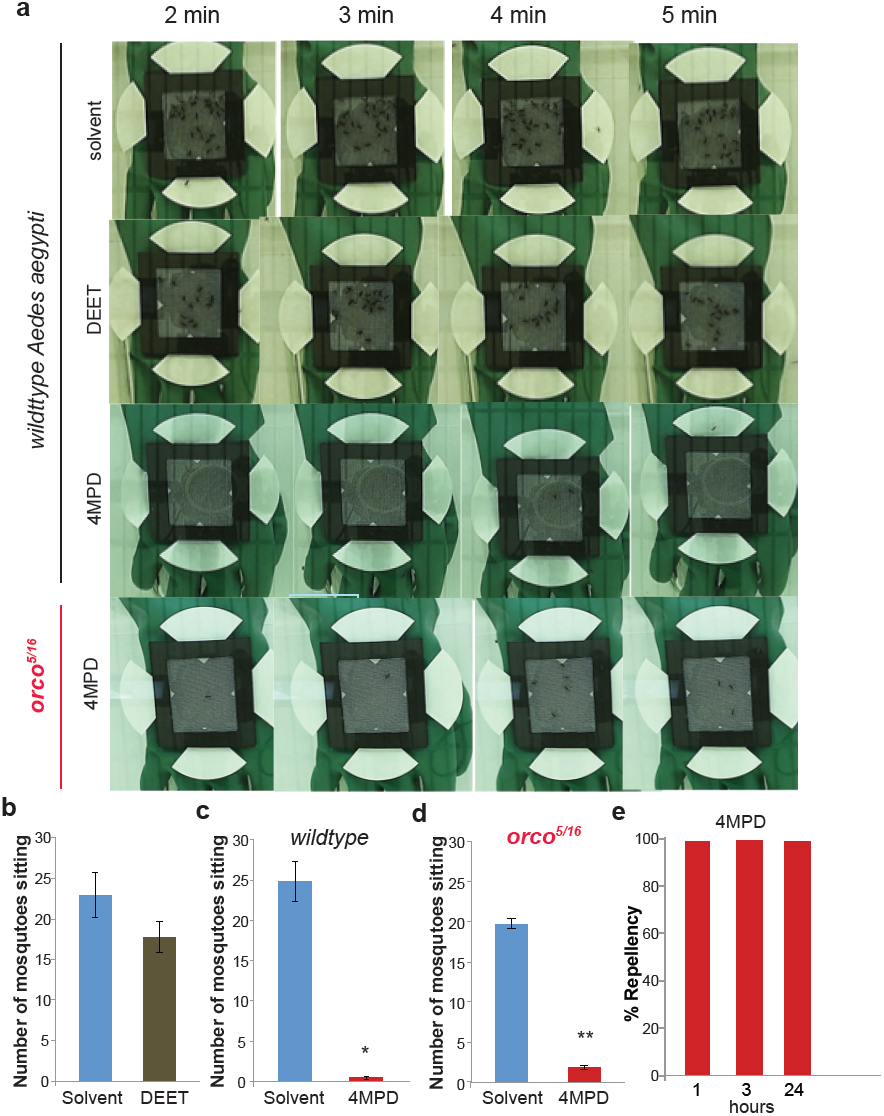
Higher volatility 4-methylpiperidine repels more spatially than DEET, and does not depend on *Or* family receptors. **a,** Representative images of the spatial arm-in-cage behavior assay from minutes 2,3,4, and 5 for the indicated treatments in wild type and *orco*^*5/16*^ *Aedes aegypti* females. **b,c,d** Mean number of mosquito landings on the net in the solvent and indicated treatments at 3%. N=5 trials, ∼40 mosquitoes/trial. * = p<0.005, ** = p< 0.001. **e,** Percentage repellency measured at 1 hr, 3 hr and 24 hr time points after application of the compounds to the test net (stored at 37C) in a standard arm-in-cage assay for 4-MPD (10% conc.). N=3 trials each, 24 hr=1 trial, ∼40 mosquitoes/trial.

### 4-methylpiperidine repellency is not mediated by the odorant receptor gene family

Analysis of olfactory (non-contact) repellency of DEET is often confounded by gustatory (contact) avoidance in behavioral assays, making it difficult to parse out the contribution of olfactory avoidance. Since 4-methylpiperidine has a spatial repellent effect, testing the contribution of the olfactory system is experimentally straightforward. We tested *orco* co-receptor mutant *A. aegypti* in which all Ors are rendered non-functional, in the spatial arm-in-cage assay. *orco* mutant *A. aegypti* mosquitoes avoided 4-methypiperidine very strongly, demonstrating that *Or* family receptors are not required for avoidance of this compound (Fig 3d). This observation is consistent with our finding that 4-methypiperidine is an Ir40a+ neuron activator, and our model that, conserved *Ir* gene family members, and not those of the *Or* family, detect the repellent.

Volatile repellents can have limited time of protection. In experiments using a standard arm- in-cage protocol, we tested the duration of repellency for 4-methylpiperidine over a 24 hr period. The 4-methylpiperidine (10%) showed a100% repellency across all 3 time-points tested: 1 hr, 3hr and 24hr (Fig 3e). Taken together these results suggest that a repellent that activates *Ir40a* neurons can protect a spatial zone, and can last more than 24 hrs.

### Orco- mosquitoes retain avoidance to DEET

Our finding that repellency to 4-methypiperidine in *A. aegypti* does not require *orco* prompted us to take a second look at another agonist of *Ir40a* neurons, DEET. A previous study reported that *orco* mutant *A. aegypti* lose olfactory repellency to DEET (DeGennaro, McBride et al. 2013). However, the same study also reported that *orco* mutant *A. aegypti* do not in fact bite a DEET covered arm, an effect that was considered to rely solely on contact repellency via gustatory neurons. Our previous findings suggested that contact avoidance plays a minor role in the overall repellency to DEET in *Aedes* mosquitoes (Kain, Boyle et al. 2013). Moreover, DEET has been shown to have a fixative effect that dramatically alters the composition of volatiles emitted from skin (Syed and Leal 2008), which may confound interpretation of results from experiments in which DEET was applied directly onto a human arm and placed near to a cage of mosquitoes (DeGennaro, McBride et al. 2013). We therefore decided to test non-contact DEET repellency in *orco* mutant mosquitoes in 2-choice assays that use heat as a simple non-olfactory attractant (Fig 4a), which removes confounds of volatile fixative effects and inter-person variations in skin chemistry. Wild type *A. aegypti* females avoided the DEET treatment side as expected (Fig 4b,c). Surprisingly, *orco* mutant females also showed robust avoidance of the DEET-treated side (Fig 4b,d). These results indicate that even in the absence of any *Or-*mediated repellency of DEET, robust olfactory avoidance of DEET occurs. This finding is consistent with our model that an Or-independent Ir pathway also mediates strong DEET avoidance.

**Figure 4.**
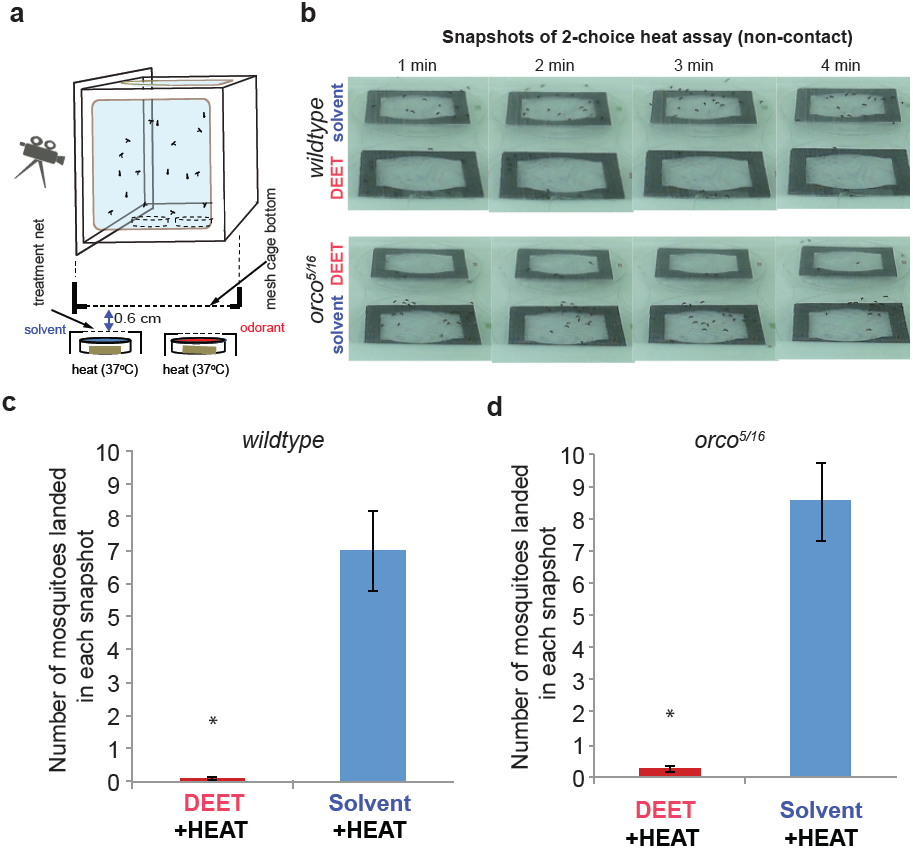
*Aedes aegypti* mosquitoes mutant for *orco* are still strongly repelled by DEET from an attractive heat source or human arm. **a,** Schematic of the 2-choice heat attraction assay where two Petri dishes each with a ∼37C heat source and a net-covering which is treated with test chemicals, is placed 6 mm below a cage containing ∼40 mosquitoes. **b,** Representative images of 2-choice heat attraction assays in *Aedes aegypti* females of *wildtype* and *orco*^*5/16*^ genotype to DEET (3%), and **b,c,** Mean number of mosquitoes of the indicated genotype present on the untreated net interior of cage at any timepoint (measured at times 2,3,4,5 min) above solvent treated or DEET treated heat pads. N= 10 trials, ∼40 mosquitoes /trial. *=p<0.005.

## Discussion

By performing activity screening on *Ir40a-*neurons we were able to identify a strong agonist that acts as a repellent for both flies and mosquitoes. The identified molecule does not share structural features with DEET, and has higher volatility, demonstrating the utility of our neuronal-activity based approach in finding novel repellents. While 4-methylpiperidine is not appropriate for human use due to its safety and smell characteristics, these proof-of-principle experiments demonstrate that we will be able to use the *Ir40a* neuron to identify effective repellents that possess a variety of different physicochemical properties.

Such methods could enable development of completely new chemical classes of repellents that could protect people over extended spatial ranges or over prolonged periods of time. The more classes of volatile repellents are discovered, the greater choice there will be in terms of important usability parameters such as vapor pressure, smell, safety, and solubility, thus expanding possibilities for use in different formulations such as body lotions, skin creams, washing detergents, perfumes, as well as application on clothes, bednets, house entryways, backyards, candles, and evaporators.

Since the *Ir* receptors are highly conserved across insect species, new generations of repellents may also be used to reduce disease transmission by other vectors that cause severe morbidity, mortality, and economical losses such as Tse Tse and Sand flies. Furthermore, these repellents may be used against insects that damage crops and enter homes, since the genomes of most agricultural pests and urban pests encode the conserved *Ir40a, Ir93a* and *Ir25a* genes. Importantly, new repellents that are structurally dissimilar from DEET could potentially be effective against DEET-resistant strains. *Ir40a*-expressing neurons also express two additional receptors, *Ir25a* and *Ir93a.* The exact subunit structure of the receptor is still unknown, but given the structural diversity of the ligands, it is possible that more than one binding site may be present.

Recent studies have implicated two distinct pathways in the detection of DEET and its analogs in the insect olfactory system: the *Ionotropic receptors* that are ancient and highly conserved, and the *Odorant receptors* that are diverse and poorly conserved. The insect olfactory system is extremely sophisticated, with receptors from these two pathways contributing to detection of volatiles in numerous different olfactory neurons. More often than not, single compounds are detected combinatorially by detection with multiple receptors. It is conceivable, and almost expected, that repellents are also detected by multiple receptors, as we indicated in a model proposed at the end of a recent paper (Kain, Boyle et al. 2013). Correspondingly, an analysis of the literature reveals a few reports of *Or* family receptors identified from *ex vivo* cell-based assays as being either activated by DEET, or inhibited (Liu, Pitts et al. 2010; Pellegrino, Steinbach et al. 2011; Bohbot and Dickens 2012; Xu, Choo et al. 2014).

Amongst these, the role of an *Or* family member, *CqiOr136*, in DEET avoidance was proposed in *Culex quinquifasciatus* (Xu, Choo et al. 2014). However, orthologs of *CqiOr136* are not present in *A. aegypti, D. melanogaster* or any other important vector including *Anopheles gambiae* and its related species, *Phlebotomous* flies, and *Glossna morsitans*. This apparent incongruence with evolutionary conservation of DEET’s repellency is further complicated by the fact that some arthropods that are repelled by DEET, such as ticks, do not encode the *Or* family in their genomes. Furthermore, the conclusion that *CqiOr136* is required for DEET repellency arises from behavior assays using an irrelevant low concentration of DEET, at 0.1%, whereas formulations for human use are usually 100-1000 fold higher, raising concerns that other receptors may the most important contributors at relevant concentrations.

The role of the *Or* family in DEET repellency of mosquitoes therefore rests largely on the results of a study in which *orco* mutant *A. aegypti* were shown to be attracted to a region of a cage placed adjacent to a human arm covered with DEET (DeGennaro, McBride et al. 2013). However, the study also reported that *orco* mosquitoes did not bite the DEET-covered arm, and video analysis used by the authors suggested that the lack of bites was a consequence of gustatory contact avoidance. If olfactory repellency were entirely lost in *orco* mutants, it would largely invalidate our findings that *Ir* receptors are also major contributors to olfactory avoidance of DEET. A human arm is a complex mixture of hundreds of odorants, heat and humidity, which together comprise highly attractive cues that are extremely variable from person to person. Moreover, direct application of DEET is known to have a fixative effect that significantly alters skin odor emissions (Syed and Leal 2008). In taking a reductionist approach to understanding the role of *Or* pathways, we use a simple assay using heat as a non-olfactory attraction cue and now find that *orco* mosquitoes indeed retain substantial aversion to volatile DEET. The weaker attractant cue (heat) likely enables us to understand repellency in the mosquitoes lacking *Or* function completely. This result unambiguously demonstrates that mechanisms other than *Or* receptors also play a role in repellency to DEET. The role of non-*Or* family receptors in aversion to 4-methylpiperidine is also unequivocally revealed in the arm-in-cage assays. Together these findings are consistent with our model that *Ionotropic receptors* such as *Ir40a* play an important role in aversion. We cannot rule out the possibility that additional receptors, *Irs* or otherwise, may also contribute to repellency, given that *Ir* receptors can have overlapping ligand responses and DEET avoidance is so well conserved across insects. Further studies of the *Ir* receptor repertoire, as well as other non-canonical candidate receptors, in mosquitoes will be required to completely describe the underlying mechanisms of DEET repellency and to find promising new targets for identifying safe and effective repellents.

Overall, our findings validate the activity of the *Drosophila Ir40a* neuron as a predictor of repellency, and demonstrate that behavioral testing in the model insect *Drosophila melanogaster* is an efficient means to identify new generations of insect repellents that can have broader activity against more dangerous vectors like mosquitoes. These findings set the stage for identification of new generations of broad-spectrum insect repellents that have enhanced activities and desirable properties, and provide a foundation for discovery of superior repellents in the future.

## Methods

### Fly stocks

The *UAS-GCaMP3* was obtained from Bloomington Stock Center (BL#32236) and the *Ir40a-GAL4* stocks from R. Benton. Fly stocks were grown on cornmeal-dextrose media, at 25°C unless otherwise noted.

### Calcium imaging

Flies of genotype *Ir40aGAL4/CyO; UAS-GCaMP3* were used for calcium imaging after they were 2 weeks old essentially as described earlier (Kain, Boyle et al. 2013). A fly was immobilized inside a 200 ul yellow pipette tip (tip cut back) with their antennae protruding out. One antenna was held down using a glass electrode on a thin layer of 70% glycerol that enhanced imaging of fluorescence. The antenna was orientated with the arista and sacculus pointing upwards, which is important for accessibility to odor stimuli. Odorants in water or paraffin oil (100ul) were applied onto 2 Whatman filter paper strips (2x3 cm) placed inside 5ml plastic syringes, and delivered manually over the preparation. Fresh solutions were prepared each time for a 20% 4-methylpiperidine solution in distilled water, 10% Ethyl acetate and 10% Butanol solutions in paraffin oil. GCaMP3 fluorescence intensity in the neurons within the sacculus of the antenna was visualized with a Leica SP2 confocal microscope with a 20X air objective. Samples were excited with a 488-nm laser, and emissions were collected through a 505–530 band-pass filter. Images were acquired at one frame every 3 s at a resolution of 512 X 512 pixels and 50–100 frames were acquired for each odor application, starting at least 10 frames before application of the stimulus and 40–90 frames during and after stimulus application. Odor puffs (∼ 2 sec) were delivered manually over the antenna by from the 5ml plastic syringe. Leica SP2 software and ImageJ were used to calculate minimum and maximum fluorescence intensities in labeled cell bodies for each stimulus application, which was used to calculate **Δ** F/F% as earlier (Kain, Boyle et al. 2013).

### Drosophila Behavior

Wild-type *Drosophila melanogaster* (w^1118^) 20 male/20 females per trial were starved for 24 hrs and tested when 3-7 days old in the t-maze assay as in (Kain,Boyle et al. 2013).

### Mosquito Behavior

Mosquito behavior assays were done with 40 mated, non-blood fed and 24hr starved, 5-10 day old females in 30cm^3^ cages fitted with clear glass tops. *Aedes aegypti* Orlando strain and *orco*^*5/16*^ mutants (provided by L. Vosshall) were used in behavior experiments.

#### Arm-in-cage assays (time course)

Mosquito arm-in-cage assays were performed as previously described (Kain, Boyle et al. 2013) in a contact repellency setting with a few modifications. A set of five 1.5mm thick magnetic window frames cut from flexible magnet strips (McMaster-carr #4 77K24) were used to secure a treated net over the glove window. Solvent treated and odorant treated nettings were hung in fume hood until dry and then assembled for testing. Treatment and control nettings were tested, then stored at ambient conditions and assayed after 1hr, 3hrs and 24 hours interval.

#### Modified arm-in-cage spatial repellency

A rectangular 7 X 6 cm piece of untreated net was used on top of the glove window in this assay positioned 6 mm above skin surface. A circular piece of filter paper 90mm diameter (Whatman #1) was cut into quadrants. One piece was secured at the pointed edge on each side of the glove window by a magnet layer below, but not in contact with the net. Test 4-methylpiperidine solution (100μl) of 3% concentration was added evenly to each filter paper piece. A hand was inserted into the glove and placed inside the test cage for 5 minutes. Each cage of mosquitoes was only used for one trial.

#### Two-choice heat assay (non-contact)

A pair of heat sources, for test and control was prepared using 4 hand warmers (Hot hands® Hand warmers HH2; Heat Max, Dalton, GA) that were simultaneously activated by shaking in gloved hands to 37°C. The hand warmers were fitted into a 100 x 15 mm Petri dish base and covered with 15 x 15 cm polyester netting secured round the Petri dish by a pair or 8 inch plastic cable ties (Gardner Bender, Milwaukee, WI) coupling. Excess netting material was trimmed off round the edges of the Petri dish. These heat source petri dishes were each covered with a 150 X 15 mm petri dish base with a 10 x 7.5 cm window opening cut out of it. The window opening was then aligned with the top of the heat source petri dish underneath and secured in position with the aid of the loose ends of the cable ties. The pair of large petri dish assembly was placed side by side with the window openings aligned to position. Another pair of 150 x 15mm petri dish bases was placed next to the assembly to keep the arena in position and to act as support base for test cage. Pieces of 9 x 8 cm nets treated with (500μl) solvent or DEET 3% concentrations were suspended in a fume hood to allow solvent evaporation. Each treatment piece was placed between two 10 x 7.5 cm flexible magnets with 7.5 x 6.2 cm window frame. Three magnetic window frames were added on top. These solvent and odorant (DEET) treatment net- magnets assembly were simultaneously placed directly over the 10 x 7.5 cm window openings of the larger petri dishes on top the heat pads forming a 2 – choice assay arena. At the start of the assay, each test cage was gently set on its side aligned directly over the 2-choice arena. The pair of petri dish bases and the magnets layers always maintained a gap of 6 mm between the lower treated net and the test mosquito cage screen side thus ensuring no contact between mosquitoes and the treatments. Mosquitoes attracted by heat were exposed to either DEET or acetone solvent treatment in a non-contact manner. The solvent and DEET positions were alternated between runs. Mosquito landing choices were recorded during the assay and videos manually analyzed by counting the number of landed mosquitoes in snapshots taken at 1 minute interval from the second minute for the duration of the 5 minute trial.

#### Two-choice heat assay (contact)

A pair of heat source petri dishes were prepared as before using activated hand warmers fitted into 100 x 15 mm petri dish bases. The treatment chemical (500μl) was added at 3% concentration directly onto the 15 x 15 cm polyester net and then hung to dry to. The net was secured round the petri dish heat source by a pair of 8 inch plastic cable ties) coupling. Excess netting material was trimmed off round the edges of the petri dishes. At start of the assay, the two assembled dishes were placed side by side inside a test cage with mosquitoes and mosquito behavior recorded for 5 minutes. Total numbers of mosquito landings on the net covering of each dish were counted manually from video recordings for the 5 min duration of the assay. The solvent and DEET positions in the test cages were alternated between runs.

## Acknowledgements

We would like to thank Greg Pask and Anupama Dahanukar for comments on the manuscript. This work was funded in part by the UCR NIFA funds. Funder had no role in experimental design or writing.

